# Loss of primary cilia and dopaminergic neuroprotection in pathogenic LRRK2-driven and idiopathic Parkinson’s disease

**DOI:** 10.1101/2024.01.15.575737

**Authors:** Shahzad S. Khan, Ebsy Jaimon, Yu-En Lin, Jonas Nikoloff, Francesca Tonelli, Dario R. Alessi, Suzanne R. Pfeffer

## Abstract

Activating LRRK2 mutations cause Parkinson’s disease. Previously, we showed that cholinergic interneurons and astrocytes but not medium spiny neurons of the dorsal striatum lose primary cilia in LRRK2 mutant mice. Single nucleus RNA sequencing shows that cilia loss in cholinergic interneurons correlates with higher LRRK2 expression and decreased glial derived neurotrophic factor transcription. Nevertheless, much higher LRRK2 expression is seen in medium spiny neurons that have normal cilia in mice and humans. In parallel with decreased striatal dopaminergic neurite density, LRRK2 G2019S neurons show increased autism-linked CNTN5 adhesion protein expression; glial cells show significant loss of ferritin heavy chain. Human striatal tissue from LRRK2 pathway mutation carriers and idiopathic Parkinson’s disease show similar cilia loss in cholinergic interneurons and astrocytes and overall loss of such neurons. These data strongly suggest that loss of cilia in specific striatal cell types decreases neuroprotection for dopamine neurons in mice and human Parkinson’s disease.

**Teaser:** Cilia loss in Parkinson’s disease decreases dopaminergic neuroprotection due to inability to sense Hedgehog signals

## Introduction

Mutations in the gene encoding leucine rich repeat kinase 2 (LRRK2) cause inherited Parkinson’s disease (PD), a neurodegenerative disorder that results in loss of dopaminergic neurons in the Substantia nigra (*1–4*). LRRK2 phosphorylates a subset of Rab GTPases (*2, 3*), and reversal of LRRK2 phosphorylation is mediated at least in part by the PPM1H phosphatase (*5*). LRRK2 action blocks the process of primary cilia formation (*6, 7*) and loss of PPM1H in wild-type mouse embryonic fibroblast cells phenocopies the loss of cilia seen upon expression of pathogenic LRRK2 (*5*).

Rab GTPases are master regulators of protein trafficking and carry out their roles by binding to specific partner proteins when GTP-bound (*8, 9*). Phosphorylation of Rab proteins interferes with their abilities to be loaded with GTP by guanine nucleotide exchange factors, a prerequisite for their binding to partner effector proteins (*2, 3*). Instead, once phosphorylated, Rab GTPases bind to new sets of phospho-specific protein effectors. For Rab8 and Rab10 these include RILPL1, RILPL2, JIP3, JIP4, and Myosin Va proteins (*3, 6, 10, 11*). These phospho-Rab interactions block ciliogenesis in cell culture and mouse brain via a process that requires RILPL1 and Rab10 proteins (*3, 6, 7*); centriolar cohesion is also altered (*12, 13*).

The dorsal striatum is composed primarily of medium spiny neurons, interneurons and glial cells such as astrocytes, and this region is infiltrated by the extensive processes of dopamine secreting neurons from the Substantia nigra (*14*). Nigral dopaminergic neurons secrete Sonic Hedgehog (Shh) that is sensed by poorly abundant, striatal cholinergic interneurons (*15*). Shh is needed for the survival of both the cholinergic target cells and the Shh-producing dopaminergic neurons, even though only the cholinergic neurons express the PTCH1 Shh receptor. Shh triggers secretion of glial derived neurotrophic factor (GDNF) from the cholinergic neurons, which provides reciprocal neuroprotection for the dopaminergic neurons of the Substantia nigra (*15*).

We showed previously that the rare, striatal, cholinergic interneurons that would normally sense Shh via their primary cilia are significantly less ciliated in four different mouse models of pathogenic LRRK2 disease (LRRK2 G2019S knock in or transgene, R1441C knock in, or PPM1H knockout (*6, 16*). In that work, we predicted that cilia loss would decrease the ability of cells to sense Shh signals and we found strong evidence of Shh signaling dysfunction due to cilia loss in cholinergic neurons as well as astrocytes that share a broad ciliary deficit. Here we report single nucleus RNA sequencing of the dorsal striatum of wild type and G2019S LRRK2 mutant mice. We see limited overall transcriptional changes, decreased GDNF transcripts in cilia-deficient LRRK2 G2019S cholinergic neurons, increases in the RNA encoding the autism-spectrum disorder-associated Contactin 5 neuronal adhesion protein and down regulation of the iron-binding ferritin heavy chain mRNA. Remarkably, analogous, cell type-specific cilia loss is seen in human postmortem striatum from patients with both idiopathic and LRRK2-pathway based PD.

## RESULTS

In an attempt to determine why cilia are only lost in certain cell types in the striatum of mice harboring activating LRRK2 mutations or missing the PPM1H phosphatase, we carried out single nucleus RNA sequencing (snRNAseq) of dorsal striatum from 6-month-old, wild type and LRRK2 G2019S mice. Shown in Figure 1A is a t-distributed Stochastic Neighbor Embedding (tSNE) plot used to identify distinct cell types in this brain region of wild type mice. These clusters represent 54,023 nuclei that were sequenced from 6 female mice. As expected, the largest clusters represent the direct and indirect spiny neurons that express D1 and D2 dopamine receptors respectively, followed by oligodendrocytes, astrocytes and microglia. Much smaller clusters were detected representing neuronal precursor cells, eccentric spiny neurons, ependymal, endothelial and mural cells, as well as parvalbumin, cholinergic and somatostatin-expressing interneurons (Fig. 1A). Clusters were identified using established markers of each cell type in this brain region (Table 1) (*17–22*); clustering was based on 14 cell types in this representation.

**Figure 1.**
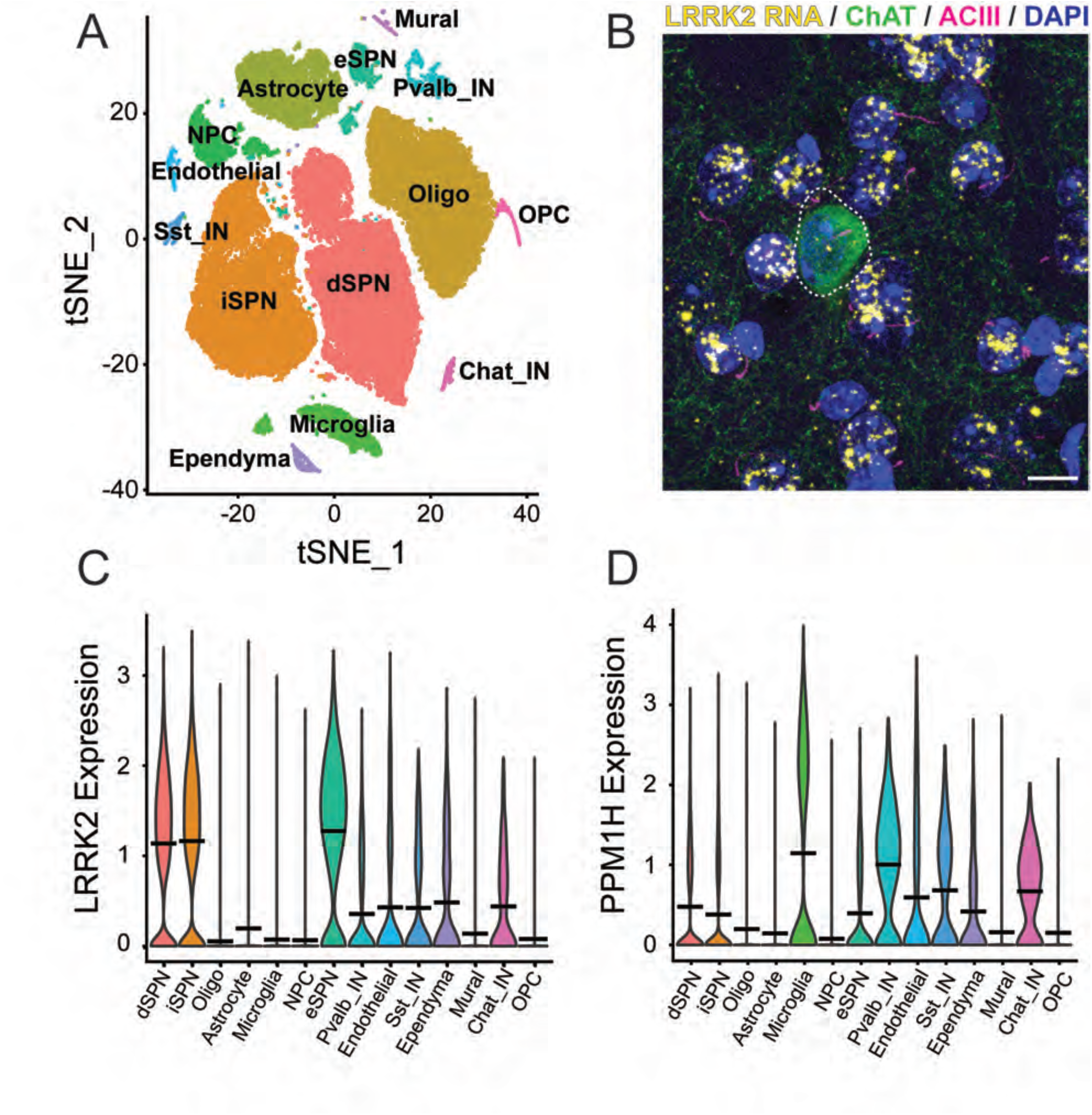
Single nucleus RNA sequencing analysis of wild type mouse dorsal striatum. A. tSNE plot showing cell types detected using markers summarized in Table 1. C, D. Comparison of relative LRRK2 (C) or PPM1H (D) RNA levels in cell types color coded as in A. B. RNA scope in situ hybridization to detect LRRK2 transcripts in the dorsal striatum. Green, anti-choline acetyltransferase staining of a cholinergic interneuron surrounded by the much more abundant medium spiny neurons; pink, primary cilia; yellow dots, LRRK2 transcripts, blue, DAPI stained nuclei. Bar, 10µm.

**Table 1.** Markers used to identify cell type clusters.

| Cluster ID | Key Markers |
| --- | --- |
| **dSPN** Direct spiny neuron | Drd1, Lingo2 |
| **iSPN** Indirect spiny neuron | Drd2, Adora2a |
| **Oligo** oligodendrocyte | Stl18, Pex5l |
| **Astrocyte** | Gpc5, Slc1a2 |
| **Microglia** | Cx3cr1, Inpp5d |
| **NPC** Neuronal precursor cell | Sox4, Dlx1 |
| **OPC** Oligodendrocyte precursor | Tnr, Pdgfra |
| **eSPN** Eccentric spiny neuron | Otof, Adarb2 |
| **Pvalb_IN** Parvalbumin interneuron | Kit, Nxph1, Ilrap2 |
| **Endothelial** | Flt1, Slcola4 |
| **Ependymal** | Dnah12, Tmem112 |
| **Mural** | Atp13a5 |
| **Sst_IN** Somatostatin interneuron | Nos1 |
| **Chat_IN** Cholinergic interneuron | Cpne4, Clstn2 |
| **Fibroblast** | Xfp804b |

One reason why astrocytes and cholinergic neurons, but not the much more abundant medium spiny neurons lose their cilia might be due to their relative content of LRRK2 kinase or the counteracting, PPM1H phosphatase. SnRNAseq showed that at least at the RNA level, the reverse is true: direct comparison of RNA transcript abundance showed that direct, indirect and eccentric spiny neurons express the highest relative amounts of LRRK2 RNA compared with astrocytes and cholinergic interneurons (Fig. 1C). Microglia, parvalbumin and cholinergic interneurons also expressed slightly higher levels of PPM1H than other cell types (Fig. 1D). RNAscope fluorescence in situ hybridization confirmed this discrepancy as shown in Fig. 1B; the predominant, ciliated medium spiny neurons showed many more LRRK2 transcript dots (abundant yellow spots on blue nuclei) than any of the cholinergic interneurons labeled in green with anti-Choline acetyltransferase antibody (circled with a dashed line, Fig. 1D; cilia on all neurons is shown in pink). Thus, cell-type specific, high LRRK2 levels do not necessarily predict cilia loss in the striatum.

### Two classes of dorsal striatal cholinergic interneurons

Two classes of striatal cholinergic interneurons were identified based on expression of ELAV Like RNA Binding Protein 2 (*Elavl2*) and Glutamate Metabotropic Receptor 5 (*Grm5*) (Fig. 2A left column). The *Elavl2* cluster showed lower LRRK2 expression compared with the Grm5 cluster; both clusters expressed comparable levels of PPM1H phosphatase. As reviewed by Ahmed et al. (*23*), subsets of cholinergic interneurons express either Zic4 and originate from the Septal Epithelium, or Lhx6 and originate from the Medial Ganglionic Eminence; Gbx2 is expressed in almost all cholinergic interneurons. Zic4 and Lhx6 were expressed at comparable levels between the two classes of cholinergic neurons we captured; by contrast, we did not detect Gbx2 expression.

**Figure 2.**
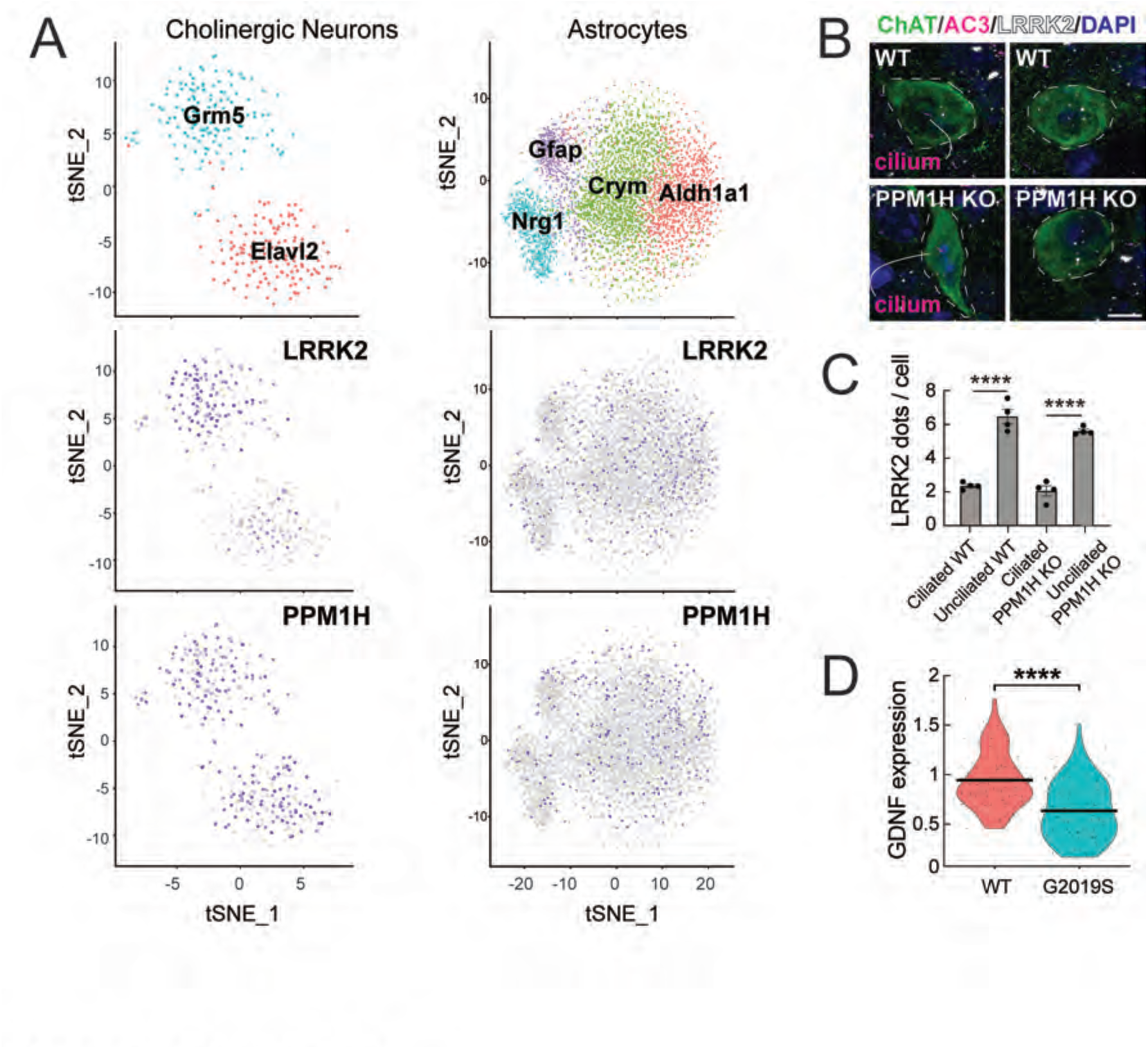
Two classes of cells explain ciliation differences in cholinergic interneurons in the dorsal striatum. A. Single nucleus RNA sequencing analysis of cholinergic interneurons (left column) or astrocytes (right column) from wild type mouse dorsal striatum. Cholinergic interneurons and astrocytes sub-clustered according to the markers indicated. Relative LRRK2 and PPM1H expression is shown below for all sub-clusters with purple intensity reflecting abundance in that nucleus. B, C. RNAscope analysis of LRRK2 RNA (white dots in B) in cholinergic interneurons according to ciliation status in 2.5-month-old wild type or PPM1H KO dorsal striatum. Error bars in C represent SEM from 4 WT and 4 PPM1H KO brains, with >25 ChAT^+^ neurons scored per brain. Statistical significance was determined using an unpaired t-test: ****p <0.0001 for ciliated WT versus unciliated WT; ****p < 0.0001 for ciliated PPM1H KO versus unciliated PPM1H KO. D. Comparison of GDNF expression from RNAseq data between WT and G2019S KI cholinergic interneurons. Statistical significance was determined using a student’s t-test. ****p <0.0001.

Because only about half of cholinergic interneurons lose their cilia in the G2019S LRRK2 dorsal striatum (*6*), we were curious to determine whether the cells that lose cilia correspond to the Grm5 subcluster that expresses higher LRRK2. Quantification of ciliation and LRRK2 RNA levels using RNAscope on dorsal striatal sections revealed that in both wild type and PPM1H knockout mice that phenocopy LRRK2 mutations (*16*), unciliated cholinergic interneurons did indeed express higher levels of LRRK2 compared with ciliated cells (Fig. 2B white dots, 2C). Thus, in cells such as cholinergic interneurons that are sensitive to LRRK2 inhibition of ciliogenesis, LRRK2 expression levels directly correlate with cilia loss. Note that wild type cells are ∼70% ciliated compared with ∼35% ciliation in PPM1H knockout striatum (*16*).

Previous work suggested that Shh signaling triggers GDNF production by striatal cholinergic interneurons (*15*). Since Hedgehog signaling is primary cilia-dependent (*24, 25*), we reasoned that loss of cilia due to LRRK2 mutation would decrease Hedgehog-responsive gene expression, including that of GDNF. Figure 2D compares the levels of GDNF RNA in cholinergic neurons of the dorsal striatum of wild type and LRRK2 G2019S mice; these neurons showed a highly significant decrease in GDNF transcription, as would be predicted from earlier findings (*15*). These data demonstrate that LRRK2 mutant mice are defective in providing critical neuroprotective factors for dopamine neurons.

### Four classes of dorsal striatal astrocytes

tSNE cluster analysis of astrocytes in the dorsal striatum revealed 4 classes of cell types based upon their relative levels of Aldehyde Dehydrogenase 1A1 (*Aldh1a1*), µ-crystallin (*Crym*), Neuregulin 1 *(Nrg1*), and GFAP (*Gfap*) RNAs (Figure 2A, right column). Chai et al., (Chai et al., 2017) noted previously that μ-crystallin displays a gradient of expression in the striatum, consistent with our findings. As reviewed by Khakh (*26*), GFAP is generally a poor marker of striatal astrocytes, while antibodies against Aldh1l1, S100β, GLT1 and Kir4.1 label most striatal astrocytes. By contrast, the astrocyte GABA transporter GAT-3 labels ∼30% of evenly distributed astrocytes. Furthermore, μ-crystallin labels ∼85% of the astrocytes in the ventral striatum, but only ∼30% in the dorsal regions, even though the density of astrocytes is equivalent. Our sequencing data are consistent with these observations in that the four astrocyte classes showed equal expression of Aldh1l1, S100β, GLT, and Kir4.1, and included both GAT-3-positive and GAT-3 negative nuclei. In addition, the four astrocyte subtypes showed comparable levels of LRRK2 and PPM1H; we do not yet know if any subcluster is more vulnerable to cilia loss than another.

### Effect of LRRK2 mutation on dorsal striatal gene expression

To further evaluate cellular changes in each of these cell types associated with the LRRK2 G2019S mutation, significant differences in RNA levels were compared for each cell type (Figs. 3, 4; Supplemental Table 1). All transcripts shown on the right side of each of the volcano plots increased in G2019S animals; transcripts shown on the left side decreased in G2019S compared with wild type mice. Figure 3 compares changes seen in direct, indirect and eccentric spiny neurons as well as Parvalbumin interneurons. The first overall conclusion from these experiments is that gene expression changes are rather minor; only an unannotated long noncoding RNA (AC149090.1) was decreased in the G2019S LRRK2 medium spiny neurons, and a larger number of transcripts increased. Nevertheless, especially noteworthy was the common increase in the RNA encoding Contactin-5 (CNTN5), an Ig-domain containing neural cell adhesion protein (also known as “NB-3” (*27*) linked to attention-deficit/hyperactivity disorder and autism spectrum disorder (*28–30*). CNTN5 interacts with CNTNAP4 as a scaffold on interneurons to support the growth of ganglion cells (*31–33*); CNTN5^-/+^ iPSC-derived glutamatergic neurons show intensified excitatory neuron synaptic activity in culture (*34*).

**Figure 3.**
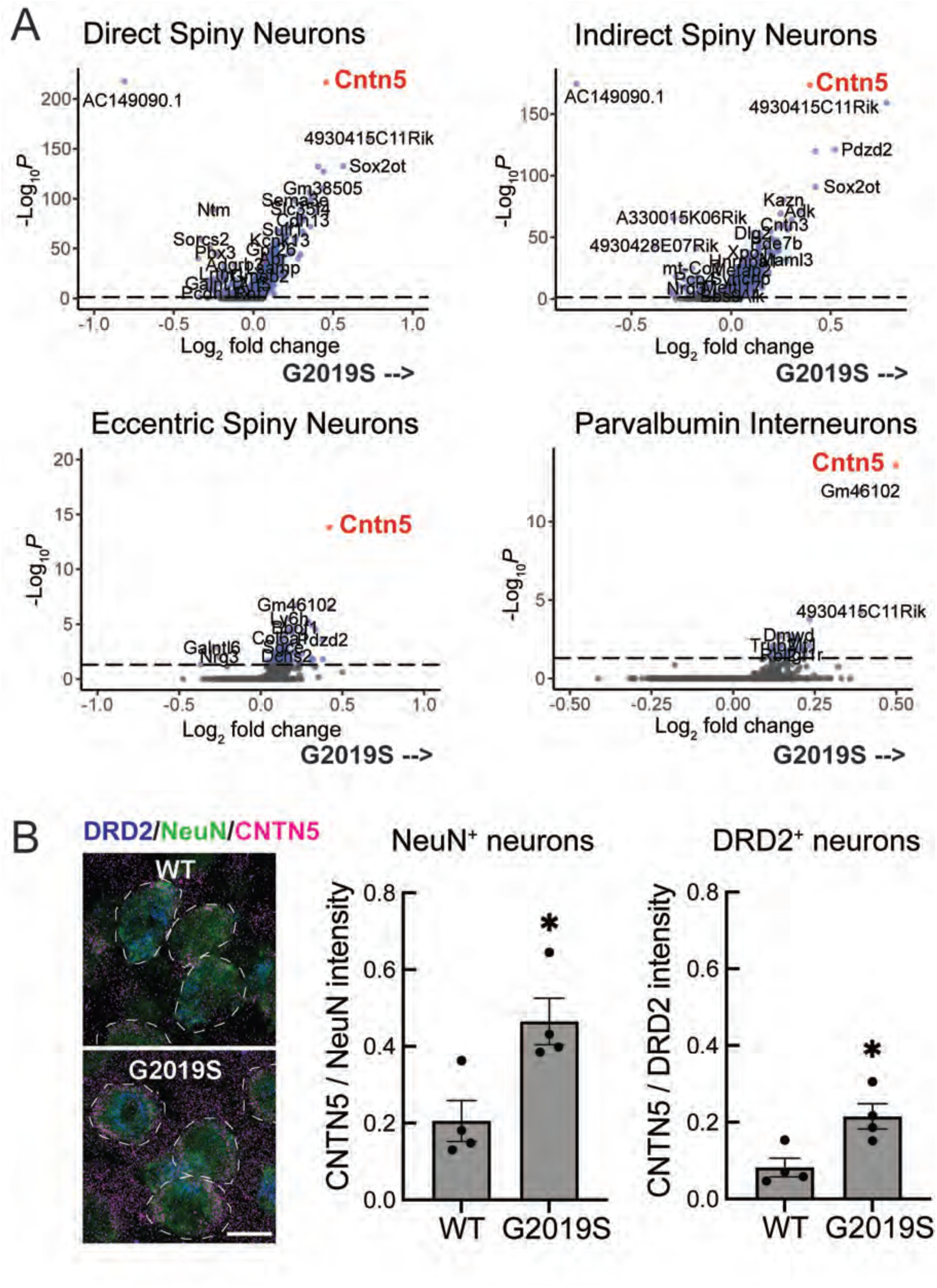
A. Volcano plot analysis comparing transcripts that increase (right side) or decrease (left side) in LRRK2 G2019S dorsal striatal neurons compared with age matched wild type mouse controls. The cell type analyzed is indicated at the top of each graph. Number of variables for direct spiny projection neurons (7699), indirect spiny projection neurons (7213), eccentric spiny projection neurons (8212), and parvalbumin interneurons (9270). B. Immunofluorescence microscopy of cells stained with antibodies to detect NeuN (green), DRD2 (blue) and CNTN5 (red) expression. Total fluorescence intensity of CNTN5 within 5 pixels of the NeuN or DRD2 was quantified and normalized to either total NeuN staining or total DRD2 staining. Error bars represent SEM from 4 WT and 4 G2019S brains, with >365 NeuN^+^ neurons and >200 DRD2^+^ neurons scored per brain. Statistical significance was determined using an unpaired t-test. CNTN5 intensity in NeuN^+^ neurons: *p = 0.0184 for WT versus G2019S and CNTN5 intensity in DRD2^+^ neurons: *p = 0.0177 for WT versus G2019S. Bar, 10µm.

Anti-CNTN5 antibody staining confirmed at least a two-fold increase in CNTN5 protein in LRRK2 G2019S NeuN-expressing neurons and DRD2-expressing indirect spiny neurons (Fig. 3B). Our working hypothesis is that loss of dopaminergic processes in the striatum of G2019S LRRK2 mice leads to upregulation of CNTN5 in spiny neurons as an attempt to strengthen or retain nigral-striatal cell-cell interactions.

### Loss of dopamine projections and GDNF receptor expression in mouse

In mice engineered such that Shh is not expressed by dopaminergic neurons, Gonzalez-Reyes et al. (*15*) reported a progressive reduction in striatal GDNF mRNA and protein, and at 12 months, upregulation of the Ret GDNF receptor and its coreceptor GDNF Receptor Alpha 1, which binds all members of the GDNF ligand family. In 5-month-old G2019S LRRK2 mice, we observed a significant ∼50% loss of striatal tyrosine hydroxylase staining compared with wild type mice, using direct quantitation of fluorescent antibody staining (Fig. 4), consistent with previous reports (*35–38*). At this stage, the level of the GFRA1 GDNF receptor present on the dopaminergic processes decreased to the same extent as tyrosine hydroxylase; no upregulation was detected (Fig. 4). The loss of striatal dopaminergic projections in G2019S LRRK2 animals is consistent with the upregulation of CNTN5 observed in the striatal spiny neurons (Fig. 3), in accord with our working hypothesis.

**Figure 4.**
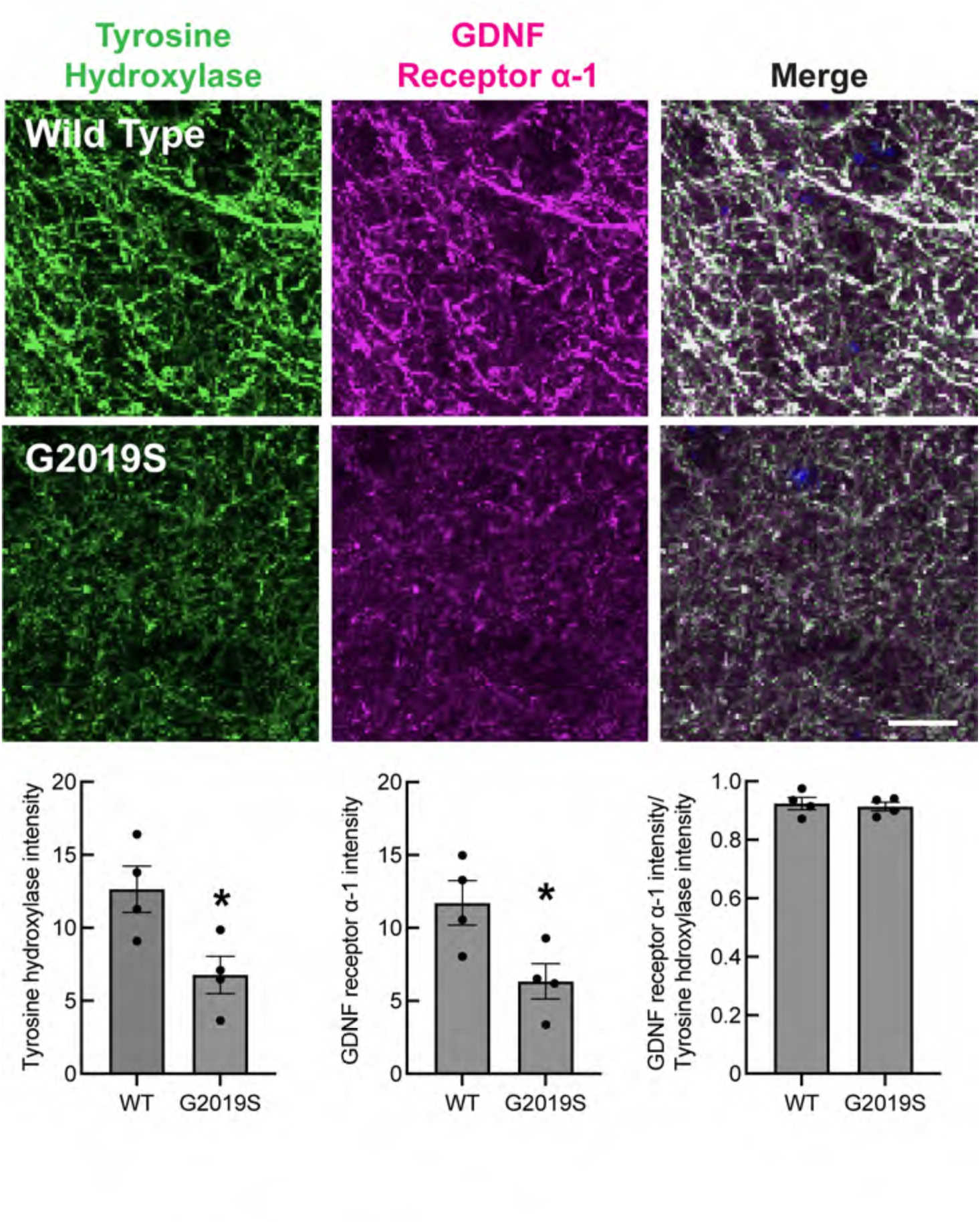
Loss of tyrosine hydroxylase and GDNF receptor staining in mouse G2019S LRRK2 dorsal striatum. Top panels: Confocal images of tissue labeled with anti-Tyrosine hydroxylase antibodies (green) and anti-GDNF receptor alpha-1 antibodies (magenta) in sections of the dorsal striatum from 5-month-old wild-type (WT) or G2019S LRRK2 KI mice. Bottom graphs: Integrated intensity of Tyrosine hydroxylase and GDNF receptor α-1 was quantified using CellProfiler. Error bars represent SEM from 4 WT and 4 G2019S brains, with >25 fields scored per brain. Statistical significance was determined using an unpaired t-test. Intensity of Tyrosine hydroxylase: *p = 0.0276 for WT versus G2019S and Intensity of GDNF receptor α-1: *p = 0.0327 for WT versus G2019S. Bar, 10µm.

### Ferritin Heavy Chain Dysregulation in LRRK2 mutant glial cells

Figure 5 shows gene expression changes seen in LRRK2 G2019S dorsal striatum compared with wild type mice for all astrocytes, oligodendrocytes, and microglia and astrocytes segregated according to individual clusters (as in Figure 2). This comparison revealed that oligodendrocytes, astrocytes and microglia all showed a decrease in Ferritin heavy chain (Fth1) transcription in G2019S LRRK2 dorsal striatum (Fig. 5). Fth1 sequesters iron within the ferritin complex, reducing the availability of free iron that can lead to generation of reactive oxygen species and lipid peroxides (*39*). A decrease in Fth1 will disrupt iron homeostasis and antioxidant defenses, increasing the likelihood of ferroptosis. Transcription of the FTH1 gene is closely tied to cellular iron levels and is normally repressed under conditions of low iron (*40*). Additional work will be needed to understand why FTH1 RNA decreases in LRRK2 G2019S oligodendrocytes, astrocytes and microglia, and whether these cells are indeed more vulnerable to ferroptosis. Low brain ferritin has been previously reported in the Substantia nigra, caudate putamen, globus pallidus, cerebral cortex, and cerebellum in human Parkinson’s disease (*41, 42*), and mouse brains deficient in ferritin heavy chain show evidence of oxidative stress (*43*).

**Figure 5.**
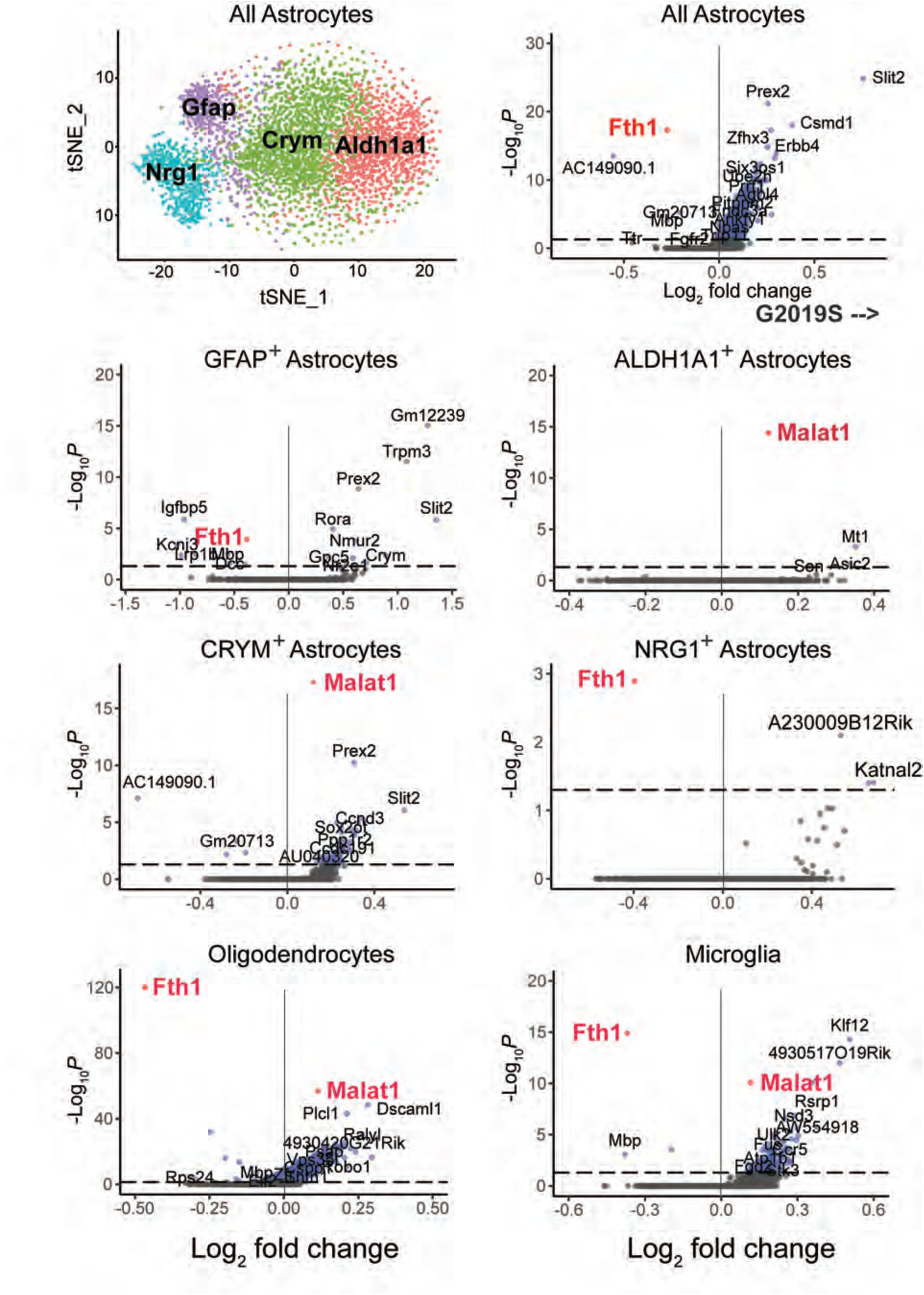
Volcano plot analysis comparing transcripts that increase (right side) or decrease (left side) in LRRK2 G2019S dorsal striatal astrocytes, oligodendrocytes or microglia compared with age matched wild type mouse controls. The cell type analyzed is indicated at the top of each graph. The tSNE plot at the upper left is the same as that presented in Fig. 2 and is included here for reference purposes. Number of genes for total astrocytes (4791), GFAP+ astrocytes (4463), ALDH1A1+ astrocytes (4901), CRYM+ astrocytes (4027), NRG1+ astrocytes (4011), Oligodendrocytes (3522 genes), Microglia (3606).

While FTH1 expression decreased in LRRK2 G2019S striatum, Malat1 RNA levels were significantly increased in CRYM^+^ and ALDH1A1^+^ astrocytes, oligodendrocytes and microglia (Fig. 5). MALAT1 is the most abundant long non-coding RNA expressed in the brain. In cultured hippocampal neurons, knock-down of Malat1 decreases synaptic density, whereas its over-expression results in a cell-autonomous increase in synaptic density (*44*). It will be interesting to determine the consequences of MALAT1 induction in these specific cell types.

### Ciliogenesis defects in human Parkinson’s disease brain

Given the importance of primary cilia in Shh signaling and the cell-type selective cilia loss observed in mouse brain, we investigated whether similar cilia loss is detected in human tissue. Table 2 summarizes the health status of the patients from which the postmortem tissue samples were obtained (*45*). As we have reported previously for mouse striatum, human striatal astrocytes and cholinergic interneurons also lose cilia in patients with LRRK2 pathway mutations as well as in patients with idiopathic PD. Figure 6A shows examples of adenylate cyclase 3 (AC3)-labeled cilia (pink) in control, human cholinergic neurons, identified using anti-choline acetyltransferase (ChAT) antibodies (green). Also shown are cholinergic neurons and their lack of cilia in striatal sections from human G2019S LRRK2 carriers, patients with sporadic Parkinson’s disease and a patient with a detrimental truncation in the PPM1H phosphatase that counteracts LRRK2 action. In mice, cilia elongate from postnatal day 7, stabilizing between P30 and P60, with ∼70% ciliation and no cilia loss up to one year of age. The control human samples from patients that were ∼85 years of age were only 12% ciliated, consistent with age-related cilia loss at later ages in humans (Fig. 6B). In contrast, all of the PD samples showed a major loss of primary cilia (Fig. 6A, B). Without cilia, these cells will not be able to sense Shh and respond by producing neurotrophic factors. Also noteworthy was our confirmation that medium spiny neurons in the human striatum retain their primary cilia, analogous to their counterparts in mouse brain: >80% of medium spiny neurons were ciliated in control and PD patient brain samples (Fig. 6C, D).

**Figure 6.**
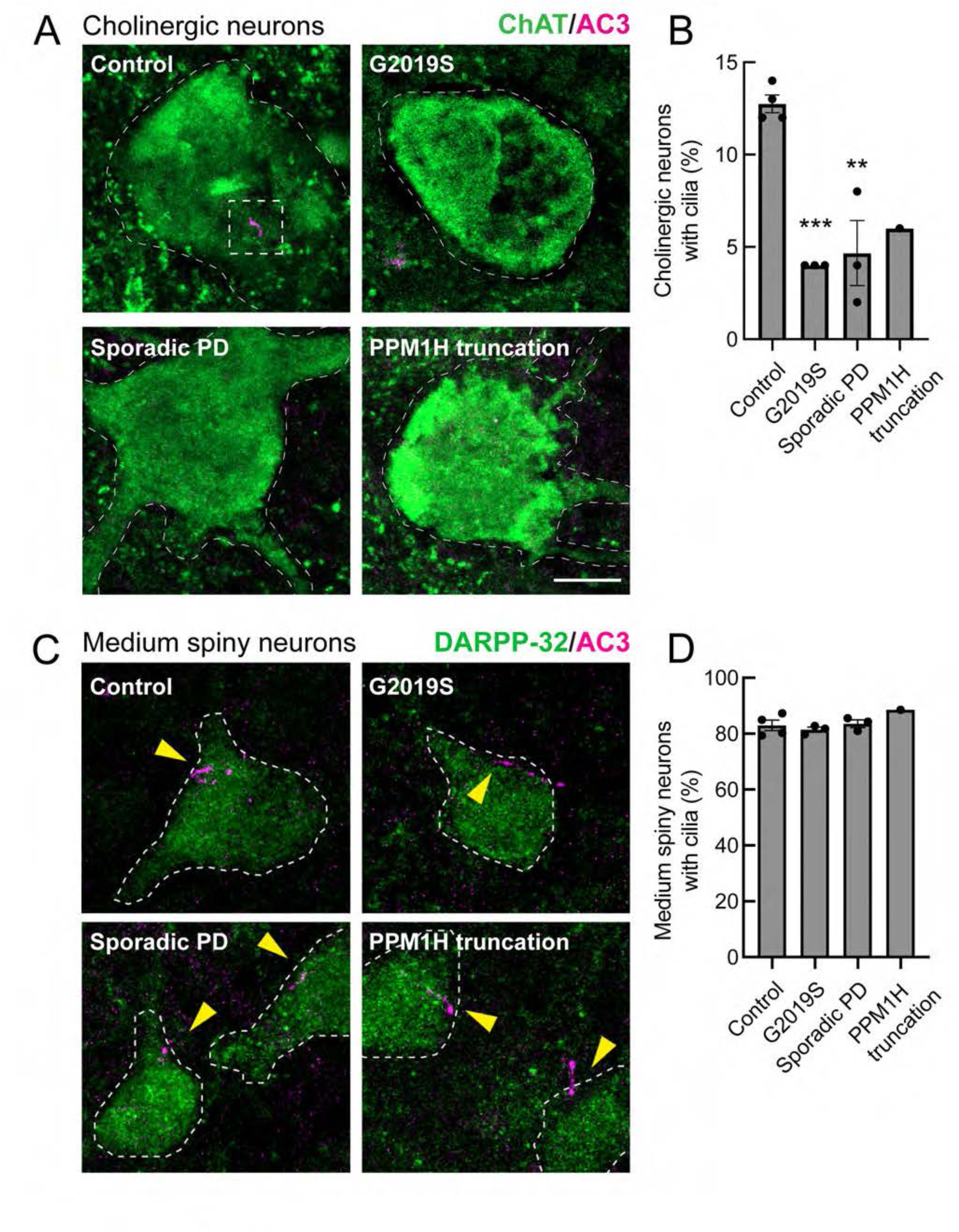
Loss of primary cilia in human striatal cholinergic neurons but not medium spiny neurons from postmortem brain sections from Parkinson’s disease patients and age matched controls. A. Confocal microscopy of tissue labeled with anti-ChAT antibody (green) and anti-AC3 antibody (magenta) to identify primary cilia. B. Quantitation of the percentage of ChAT^+^ neurons containing a cilium. Error bars represent SEM from 4 Control, 3 G2019S, 3 sporadic PD, and 1 PPM1H mutation brains, with >50 ChAT^+^ neurons scored. Statistical significance was determined using one-way ANOVA. ***p = 0.0009 for Control versus G2019S PD and **p = 0.0014 for Control versus Sporadic PD. Bar, 10µm. C. Confocal microscopy of human striatal tissue labeled with anti-DARPP-32 antibody (green) to label medium spiny neurons and anti-AC3 antibody (magenta) to identify primary cilia. D. Quantitation of the percentage of DARPP32^+^ neurons containing a cilium. Error bars represent SEM from 4 Control, 3 G2019S, 3 sporadic PD, and 1 PPM1H mutation brains, with >60 DARPP32^+^ neurons scored. All samples were scored blind and evaluated by two independent scientists. Bar, 10µm.

**Table 2.** Patient information related to human samples analyzed.

| CaseID | gender | Unified<br>Lewy Body Stage* | Summary | Age at<br>diagnosis | Age at<br>death |
| --- | --- | --- | --- | --- | --- |
| 01-31 | male | 0. No Lewy bodies | Control |  | 81 |
| 01-46 | female | 0. No Lewy bodies | Control |  | ≥90 |
| 02-27 | male | 0. No Lewy bodies | Control |  | 86 |
| 03-63 | female | 0. No Lewy bodies | Control |  | 83 |
| 10-28 | male | IV. Neocortical | Parkinson's disease | 55 | 75 |
| 10-37 | female | III. Brainstem/Limbic | Parkinson's disease;<br>Neurofibrillary tangles,<br>**LRRK2 G2019S** | 64 | 84 |
| 14-03 | male | IV. Neocortical | Parkinson's disease;<br>Alzheimer's disease; Vascular<br>dementia; **p.E309X PPM1H** | 85 | ≥90 |
| 01-39 | male | III. Brainstem/Limbic | Parkinson's disease;<br>Alzheimer's disease;<br>**LRRK2 G2019S** | 74 | 85 |
| 13-60 | male | 0. No Lewy bodies | Alzheimer's disease;<br>Parkinsonism;<br>**LRRK2 G2019S** (homozygous) | 85 | 89 |
| 16-23 | female | IV. Neocortical | Parkinson's disease | 61 | 78 |
| 18-72 | female | IV. Neocortical | Parkinson's disease | 67 | 81 |

\*Unified LB stage is the synuclein/Lewy body stage defined using the Unified Staging System for Lewy Body Disorders (Beach TG et al Acta Neuropathol.117:613-634, 2009).

Analysis of cilia in GFAP^+^ astrocytes using anti-Arl13B antibodies showed similar ciliary loss in G2091S LRRK2 patient carriers, sporadic Parkinson’s disease, and in the patient with a PPM1H truncation (Fig. 7). Control GFAP^+^ astrocytes were ∼40% ciliated while all Parkinson’s disease samples showed 10-15% ciliation. These data show that astrocyte cilia are more resistant to loss than cholinergic interneurons due to aging yet are still vulnerable to changes associated with PD.

**Figure 7.**
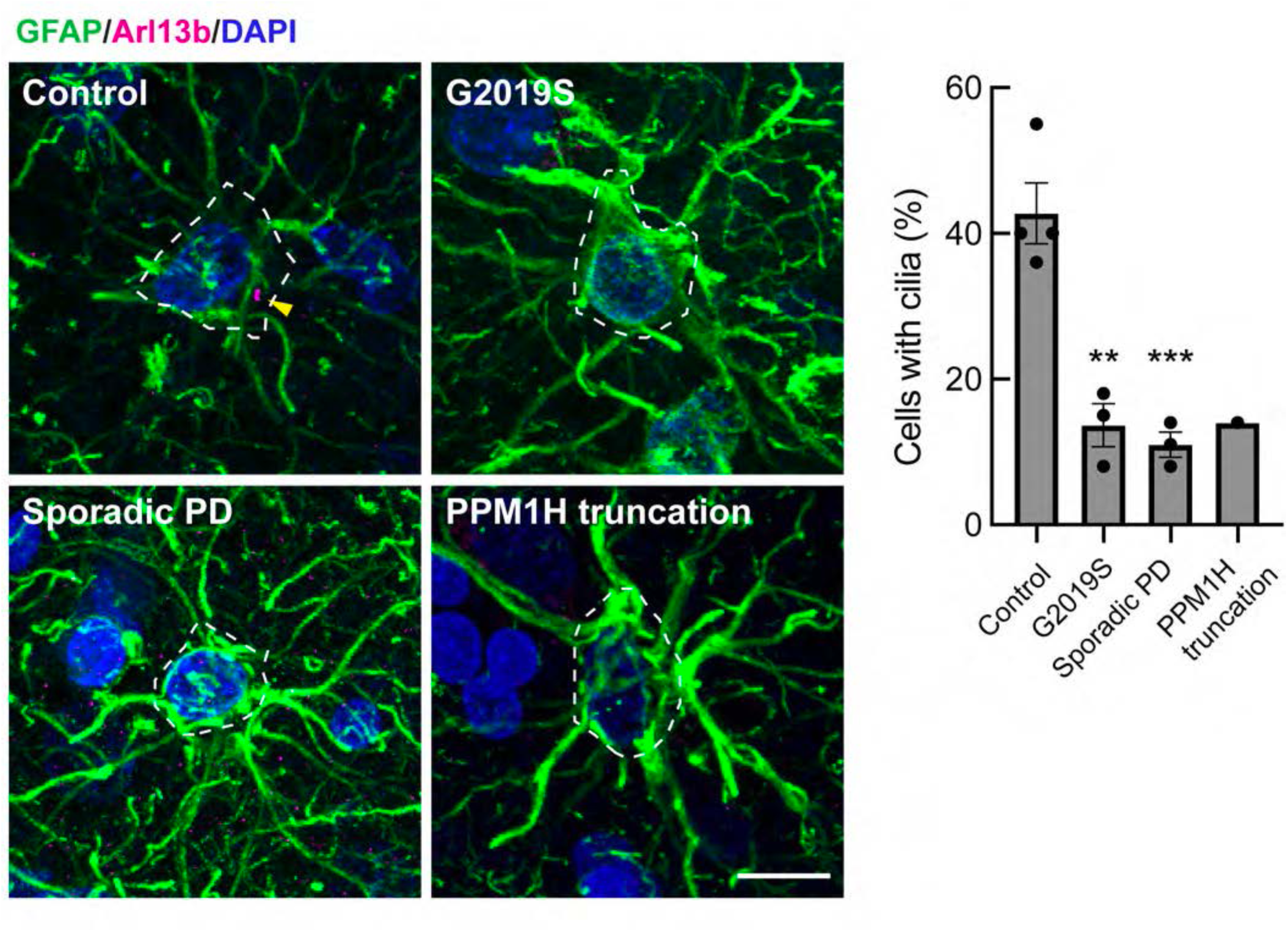
Loss of primary cilia in striatal astrocytes from postmortem brain sections from Parkinson’s disease patients. Left panels. Confocal images of astrocytes labeled using anti-GFAP (green) and primary cilia labeled using anti-Arl13b (magenta). Nuclei were labeled using DAPI (blue). Right panels. Quantitation of the percentage of GFAP^+^ astrocytes containing a cilium. Error bars represent SEM from 4 Control, 3 G2019S, 3 sporadic PD, and 1 PPM1H mutation brains, with >40 GFAP^+^ astrocytes scored. All samples were scored blind and evaluated by two independent scientists. Statistical significance was determined using one-way ANOVA. **p = 0.0015 for Control versus G2019S and ***p = 0.0009 for Control versus Sporadic PD. Bar, 10µm.

### Loss of striatal cholinergic interneurons in mice and humans

In mice lacking Shh production in dopamine neurons, progressive degeneration of cholinergic neurons was observed (*15*). In 5-month-old LRRK2 G2019S mouse striatum, we detected a trend of loss of cell number, from roughly 16 cells per mm^2^ to ∼13 cells per mm^2^ (Fig. 8A). However, in PD patient-derived brain tissue from G2019S carriers or individuals with idiopathic PD, we noted a roughly two-fold loss of cholinergic cell bodies over millimeter regions of the striatum (Fig. 8B). This is consistent with cilia loss that will decrease the ability of the interneurons to receive trophic signals including Shh from dopamine neurons or nearby Parvalbumin interneurons that also produce Shh. Because of the importance of cholinergic interneurons in coordinating the activity of striatal circuits (*46*), cell loss will also have important consequences for PD patients.

**Figure 8.**
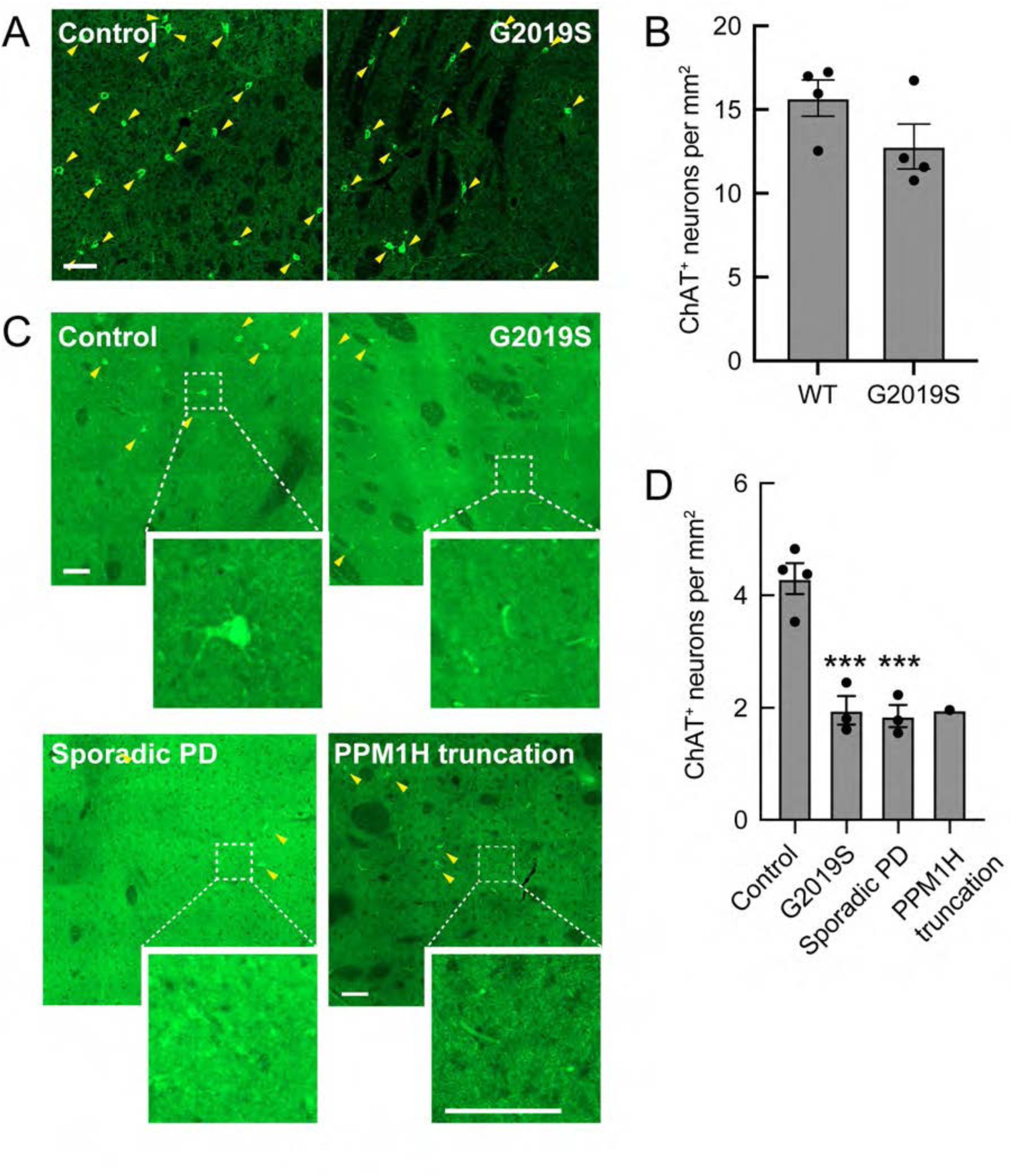
Loss of cholinergic neurons in the striatum of mice and of Parkinson’s patients. A. Representative tile scan images of the mouse dorsal striatum from 5-month-old wild-type or G2019S LRRK2 KI mice. Cholinergic interneurons were labeled using anti-ChAT (green). B. Quantitation of ChAT^+^ neurons detected per sq mm. Error bars represent SEM from 4 WT and 4 G2019S brains, with 4 sections scored per mouse. C. Representative tile scan images of the striatum from postmortem brain sections from Parkinson’s disease patients, identified geographically and molecularly using anti-DARPP-32 antibodies (cholinergic interneurons were labeled using anti-ChAT (green). D. Quantitation of ChAT^+^ neurons detected per sq mm. Error bars represent SEM from 4 Control, 3 genetic PD, 3 sporadic PD, and 1 PPM1H mutated brains, with >100 tiles scored per brain. Areas boxed with dashed lines are enlarged and shown. Statistical significance was determined using one-way ANOVA. ***p = 0.0009 for Control versus G2019S and ***p = 0.0007 for Control versus Sporadic PD. Bar, 100µm.

## Discussion

We have shown here that pathogenic LRRK2-driven cilia loss in cholinergic interneurons of the mouse dorsal striatum leads to a decrease in Shh-dependent GDNF transcription in these cells that is proportional to the extent with which they lose their cilia. Because cilia are needed for Shh signal transduction, these findings align with prior work that showed that loss of Shh expression in dopamine neurons decreases GDNF production overall in the striatum and leads to death of both cholinergic and dopaminergic neurons (*15*). We detect loss of tyrosine hydroxylase-positive dopaminergic processes in the striatum in 5-month-old G2019S LRRK2 mice, consistent with decreased availability of GDNF as a dopaminergic trophic factor; no upregulation of GDNF receptors was detected at this stage.

Prior reports have come to different conclusions as to whether LRRK2 mutant mice show dopaminergic neuron loss. Note that the experiments presented here were carried out using fluorescently tagged antibodies and at high magnification and focus on striatal processes rather than cell bodies in the Substantia nigra. Also, many prior studies used non-linear, HRP amplification methods and lower magnification and may have missed more subtle, ∼2 fold changes in tyrosine hydroxylase levels in fine processes that extend into the dorsal striatum. Under these conditions, we found that the medium spiny neurons upregulate CNTN5 transcription, perhaps as part of a last-ditch attempt to retain or stabilize inter-neuronal synaptic contacts.

Our single nucleus RNA sequencing revealed two classes of striatal cholinergic interneurons based on expression of *Elavl2* and *Grm5*. Interestingly, the *Elavl2* cluster showed lower LRRK2 expression compared with the *Grm5* cluster. ELAVL2, also called HuB, is an RNA binding protein that binds U-rich motifs in mRNAs in the 3’UTR, 5’UTR and introns of mRNAs and controls gene expression (*4, 47*). This difference in LRRK2 expression among cholinergic neurons correlated with differences in ciliogenesis: *Grm5* expressing neurons with higher LRRK2 expression were less ciliated than those expressing *Elavl2*.

Although overall gene expression changes were limited between wild type and G2019S LRRK2 mouse dorsal striatum, oligodendrocytes, astrocytes and microglia all showed a highly significant decrease in Ferritin heavy chain (Fth1) transcription in G2019S LRRK2 dorsal striatum. Oligodendrocytes are the major iron containing cells in the brain and prior work knocking out one copy of ferritin heavy chain gene rendered cells much more sensitive to oxidative stress (*48*). It is not clear why expression of this gene is particularly changed in LRRK2 mutant cells, but decreased Ferritin heavy chain will surely have major consequences for the physiology of the respective cell classes. In addition, G2019S expressing, direct-, indirect and eccentric-spiny neurons as well as parvalbumin interneurons showed increased expression of the CNTN5 neuronal adhesion protein. This parallels loss of dopaminergic process intensity in the dorsal striatum, monitored by both tyrosine hydroxylase and GDNF receptor immunofluorescence.

Analysis of postmortem human brain tissue showed low levels of astrocyte (∼40%) and cholinergic neuron (∼10%) ciliation in all samples from ∼85 year old patients, compared with ∼70% ciliation in all striatal cells of 5 month old, wild type mice. There were even fewer cilia in both LRRK2 pathway-related mutation carriers and idiopathic PD. Indeed, a patient with a deleterious truncation in PPM1H phosphatase showed the same low level of ciliation as a patient carrying the G2019S LRRK2 mutation. The lack of cilia will make these cells unable to sense Shh signals and consequently, to provide neuroprotection to their neighbors.

A remaining puzzle is why astrocyte and interneuron cilia are more sensitive to pathogenic LRRK2 expression than the predominant medium spiny neurons of the dorsal striatum in both mice and humans. Sensitivity between distinct cell type classes is not due to LRRK2 expression as our snRNAseq data showed that at least at the RNA level, LRRK2 is much more highly expressed in medium spiny neurons than in cholinergic neurons, and the counteracting PPM1H phosphatase is more highly expressed in the cells most vulnerable to LRRK2 action. Careful analyses with fully validated rabbit monoclonal antibodies have indicated that LRRK2 RNA and protein expression are concordant across the mouse brain (*49*). Future work will continue to explore the basis for the cell type specificity of LRRK2 citation inhibition.

In summary, our work reveals that specific cell types in the striatum of both mice and humans lose primary cilia due to the action of pathogenic LRRK2 kinase. Loss of cilia is seen in both LRRK2-pathway mutant carriers and in idiopathic PD, and the cause of cilia loss in idiopathic disease warrants further study. We show that loss of cilia leads to decreased GDNF production by cholinergic neurons that is needed for dopamine neuron viability. We see also loss of cholinergic neurons and dopaminergic processes, with upregulation of the neural adhesion protein, CNTN5. These data highlight the importance of neuroprotective pathways for the function of the nigrostriatal circuit.

In addition to neurons, LRRK2 G2019S astrocytes, oligodendrocytes and microglia all showed a highly significant decrease in the ferritin heavy chain, making these important cells more vulnerable to oxidative stress. Future studies using cell type specific mutant LRRK2 expression will teach us much about the relative contribution of each of these cell types to Parkinson’s disease.

## Materials and Methods

### Single nuclei RNAseq

Isolation of nuclei from adult mice was carried out as described (Allen Institute for Brain Sciences, 2021; (dx.doi.org/10.17504/protocols.io.bq7emzje) with the following modifications: Six-month old female mice (3 wildtype and 3 G2019S KI LRRK2) were trans-cardially perfused with 1x PBS. Brains were subsequently removed, dorsal striatal tissue was dissected in ice-cold PBS, placed in 1.5 mL microfuge tubes, and quickly frozen in a slurry consisting of 200 proof ethanol and dry ice. Tubes were stored at -80°C before further processing. For nuclei isolation, 1 mL of cold homogenization buffer (nuclei isolation media containing 0.1 M DTT, 1x protease inhibitor cocktail (Sigma), 0.2 U/µl of RNAsein plus and 0.1% Triton X-100) was added to each tube, and once thawed, the tissues were transferred to dounce homogenizers. Each tissue sample was dounced 10 times with a loose pestle then 10 times with a tight pestle (20 strokes total). The resulting homogenates were filtered through 30 µm strainers to remove large debris and transferred to a 15 mL conical tube. Tubes were spun at 900xg for 10 min at 4°C. Supernatants were removed, leaving only ∼50 µL of solution. Samples were resuspended in 450 µL blocking buffer (1x PBS containing 0.8% BSA and 0.2 U/µl RNAsein). 10µL of each sample was combined with 10µL of 0.4% Trypan Blue to assess nuclei quality and sample yield. Concentrations were then adjusted to 1000 nuclei/uL with blocking buffer. Sequencing using 10x Chromium Single Cell 3’, 5’, T and B cell V(D)J (V2), 10x scATACSeq, 10x 3’ WTA (V3), and feature barcoding was then performed by Stanford Genomics.

### Data Pre-processing

Seurat (version 3, https://satijalab.org/seurat/; (*50*) was used for single-cell analysis. Quality control steps were performed to identify and remove cells that were potential outliers. This included removing potential multiplets using “DoubletFinder” (https://github.com/chris-mcginnis-ucsf/DoubletFinder (*51*) and “cells” that displayed high mitochondrial gene expression (using the subset function to remove clusters with high expression of “MT-” genes). The data were then normalized and log-transformed (using the ‘LogNormalize’ method), and counts were scaled. All files, including the scripts used for Seurat gene expression analyses are available on Dryad (https://doi.org/10.5061/dryad.pk0p2ngvp).

### Analysis

Seurat (*50*) was used on the aligned dataset to identify cell clusters, and then t-SNE or uMAP was used to visualize similarities between cells. Next, cell types were assigned to these clusters based upon the expression of pre-defined marker genes (using Dropviz.org (*18*) and literature searches; see Table 1 of top hits used for cluster_IDs).

### Research standards for human and animal studies

The LRRK2^R1441C/R1441C^ mice were obtained from Jackson Laboratory (B6.Cg-*Lrrk2^tm1.1Shn^*/J mouse, JAX stock #009346 (*52*) and kept in specific pathogen-free conditions at the University of Dundee (UK). All animal studies were ethically reviewed and carried out in accordance with Animals (Scientific Procedures) Act 1986 and regulations set by the University of Dundee and the U.K. Home Office. Animal studies and breeding were approved by the University of Dundee ethical committee and performed under a U.K. Home Office project license. Mice were multiply housed at an ambient temperature (20–24°C) and humidity (45–55%) and maintained on a 12 hr light/12 hr dark cycle, with free access to food (SDS RM No. three autoclavable) and water. Genotyping of mice was performed by PCR using genomic DNA isolated from ear biopsies. Primer 1 (5’ -CTGCAGGCTACTAGATGGTCAAGGT −3’) and Primer 2 (5’ –CTAGATAGGACCGAGTGTCGCAGAG-3’) were used to detect the wild-type and knock-in alleles (<u>dx.doi.org/10.17504/protocols.io.5qpvo3xobv4o/v1</u>). Homozygous LRRK2-G2019S and littermate wild-type controls (5 months of age) were used for mouse experiments. The genotypes of the mice were confirmed by PCR on the day of experiment.

### Immunohistochemistry and microscopy

Immunohistochemistry of the human brain striatum was performed as previously described (**<u>dx.doi.org/10.17504/protocols.io.j8nlkowj1v5r/v1</u>**). Briefly, free-floating tissues containing striatum were incubated with 10 mM sodium citrate buffer pH 6.0 (preheated to 95° C) for 15 minutes at 95° C to retrieve the antigens. Tissues were permeabilized with 0.1% Triton X-100 in PBS at RT for 1 hr. Tissues were blocked with 2% FBS and 1% BSA in PBS for 2 hr at RT and were then incubated overnight at 4°C with primary antibodies. The following day, tissues were incubated with secondary antibodies at RT for 2 hr. Donkey highly cross-absorbed H + L secondary antibodies conjugated to Alexa 488, Alexa 568 or Alexa 647 were used at a 1:2000 dilution. Nuclei were stained with 0.1 µg/ml DAPI (Sigma). Tissues were incubated with freshly prepared 0.1% Sudan Black B in 70% ethanol for 20 minutes to reduce auto-fluorescence. Stained tissues were transferred to slides and overlayed with Fluoromount G and a glass coverslip. All antibody dilutions for tissue staining included 1% DMSO to help antibody penetration. All images were obtained using a Zeiss LSM 900 confocal microscope with a 63 × 1.4 oil immersion objective or 20x/0.8 objective. All image visualizations and analyses were performed using Fiji (*53*).

This table summarizes the areas analyzed for each sample:

| Brain ID | Tiles analyzed | Area analyzed (mm<sup>2</sup>) | ChAT<sup>+</sup> neurons |
| --- | --- | --- | --- |
| 01-31 (control) | 1017 | 87.60 | 309 |
| 01-46 (control) | 1082 | 91.99 | 444 |
| 02-27 (control) | 1078 | 92.63 | 413 |
| 03-63 (control) | 1041 | 82.67 | 362 |
| 01-39 (G2019S) | 645 | 55.75 | 90 |
| 10-37 (G2019S) | 117 | 10.59 | 26 |
| 13-60 (G2019S) | 637 | 54.80 | 99 |
| 10-28 (sporadic PD) | 940 | 80.36 | 143 |
| 14-03 (PPM1H) | 513 | 44.28 | 87 |
| 16-23 (sporadic PD) | 496 | 42.58 | 95 |
| 18-72 (sporadic PD) | 570 | 46.36 | 72 |

Immunostaining of the mouse brain striatum was performed as previously described (<u>dx.doi.org/10.17504/protocols.io.j8nlkowj1v5r/v1</u>): frozen slides were thawed at RT for 15 min then gently washed (2X) with PBS for 5 min. Slides were incubated with 10 mM sodium citrate buffer pH 6.0 (preheated to 95° C) for 15 minutes at 95° C to retrieve the antigens. Sections were permeabilized with 0.1% Triton X-100 in PBS at RT for 15 min. Sections were blocked with 2% FBS and 1% BSA in PBS for 2 hr at RT and were then incubated overnight at 4°C with primary antibodies. The following day, sections were incubated with secondary antibodies at RT for 2 hr. Donkey highly cross-absorbed H + L secondary antibodies conjugated to Alexa 488, Alexa 568 or Alexa 647 were used at a 1:2000 dilution. Nuclei were stained with 0.1 µg/ml DAPI (Sigma). Stained tissues were overlayed with Fluoromount G and a glass coverslip. All antibody dilutions for tissue staining included 1% DMSO to help antibody penetration. All images were obtained using a Zeiss LSM 900 confocal microscope with a 63 × 1.4 oil immersion objective. All image visualizations and analyses were performed using Fiji (*53*) and CellProfiler (*54*). RNAscope fluorescence in situ hybridization was carried out as described (*16*).

In addition to above-mentioned methods, all other statistical analysis was carried out using GraphPad Prism version 9.3.1 for Macintosh, GraphPad Software, Boston, Massachusetts USA, www.graphpad.com.

## Acknowledgments

This study was funded by the joint efforts of The Michael J. Fox Foundation for Parkinson’s Research (MJFF) (MJFF grant no. 009258 to SRP and DRA) and Aligning Science Across Parkinson’s (ASAP) initiative. MJFF administers the grant (ASAP-000463, SRP and DRA) on behalf of ASAP and itself. CYC was supported by training grant NIH 5 T32 GM007276. Funds were also provided by the Medical Research Council (grant no. MC_UU_00018/1 [DRA]), the pharmaceutical companies supporting the Division of Signal Transduction Therapy Unit Boehringer-Ingelheim, GlaxoSmithKline, Merck KGaA (DRA). For the purpose of open access, the authors have applied a CC-BY public copyright license to the Author Accepted Manuscript version arising from this submission.

We are grateful to the Banner Sun Health Research Institute Brain and Body Donation Program of Sun City, Arizona for the provision of human biological materials. The Brain and Body Donation Program has been supported by the National Institute of Neurological Disorders and Stroke (U24 NS072026 National Brain and Tissue Resource for Parkinson’s disease and Related Disorders),the National Institute on Aging (P30AG19610 and P30AG072980, Arizona Alzheimer’s Disease Center), the Arizona Department of Health Services (contract 211002, Arizona Alzheimer’s Research Center), the Arizona Biomedical Research Commission (contracts 4001, 0011, 05-901 and 1001 to the Arizona Parkinson’s Disease Consortium) and the Michael J. Fox Foundation for Parkinson’s Research.

## Author contributions

SSK, Conceptualization, investigation, formal analysis, data curation, visualization, manuscript editing

EJ and YL, investigation, formal analysis, validation, visualization, manuscript editing JN, Software, formal analysis

FT, Project administration, resources

DRA, Supervision, funding acquisition, manuscript editing

SRP, Conceptualization, writing original draft, supervision, formal analysis, funding acquisition, visualization

The authors declare that they have no competing interests. All data needed to evaluate the conclusions in the paper are present in the paper and/or the Supplementary Materials; all primary data is available at: https://doi.org/10.5061/dryad.pk0p2ngvp

---

**Table S2.** Key Resources used in this study.

| Reagent type (species) or resource | Designation | Source or reference | Identifiers | Additional information |
| --- | --- | --- | --- | --- |
| Genetic reagent(Mus musculus) | Constitutive KI Lrrk2tm4.1Arte | Taconic | #13940, RRID:IMSR_TAC:13940 | C57BL/6; G2019S KI |
| Genetic reagent(Mus musculus) | Ppm1h-/- mouse | Taconic | #TF3142 | C57BL/6 Background |
| Antibody | anti-Choline Acetyltransferase (goat polyclonal) | Millipore | AB144P-1ML (RRID:AB_2079751) | (1:200) |
| Antibody | anti-Adenylate cyclase III (mouse monoclonal) | Santa Cruz | SC-518057 (RRID:AB_3073967) | (1:100) |
| Antibody | anti-DARPP-32 (rabbit monoclonal) | Cell Signaling Technology | #2306S (RRID:AB_823479) | (1:400) |
| Antibody | anti-GFAP (chicken polyclonal) | EnCOR | CPCA-GFAP (RRID:AB_2109953) | (1:2000) |
| Antibody | anti-Arl13B (mouse monoclonal) | Neuromab | N295B/66 (RRID:AB_2877361) | (1:500) |
| Antibody | anti-GFR alpha-1/GDNF R alpha-1 (goat polyclonal) | R&D Systems | AF560 (RRID:AB_2110307) | (1:500) |
| Antibody | anti-Tyrosine hydroxylase (sheep polyclonal) | Novus Biologicals | NB300-110 (RRID:AB_10002491) | (1:500) |
| Antibody | anti-Cntn5 (rabbit polyclonal) | Novus Biologicals | NBP1-83242 (RRID:AB_11019867) | (1:50) |
| Antibody | anti-NeuN(chicken polyclonal) | Millipore | ABN91 (RRID:AB_11205760) | (1:1000) |
| Antibody | anti-DRD2(mouse monoclonal) | NeuroMab | N186/29 (RRID:AB_11000721) | (1:250) |
| Antibody | H+L Donkey anti-mouse Alexa 488 | Life Technologies | A32766 (RRID:AB_2762823) | (1:2000) |
| Antibody | H+L Donkey anti-Rabbit Alexa 568 | Life Technologies | A10042 (RRID:AB_2534017) | (1:2000) |
| Antibody | H+L Donkey anti-goat Alexa 488 | Life Technologies | A11055 (RRID:AB_2534102) | (1:2000) |
| Antibody | H+L Donkey anti-mouse Alexa 647 | Life Technologies | A31571 (RRID:AB_162542) | (1:2000) |
| Antibody | H+L Donkey anti-chicken Alexa 488 | Jackson ImmunoResearch | #703-545-155 (RRID:AB_2340375) | (1:2000) |
| Antibody | H+L Donkey anti-sheep Alexa 488 | Life Technologies | A-11015 (RRID:AB_2534082) | (1:2000) |
| Antibody | H+L Donkey anti-goat Alexa 568 | Life Technologies | A-11057 (RRID:AB_2534104) | (1:2000) |
| Reagent | Sudan Black B | Chem-Impex International | #01307 |  |
| Commercial assay or kit | RNAscopeMultiplexFluorescentReagent Kit v2 | Advanced Cell Diagnostics | #323100 |  |
| Commercial assay or kit | RNAscope Probe- Mm-Lrrk2 | Advanced Cell Diagnostics | #421551 | (1:20) |
| Commercial assay or kit | OPAL 690 REAGENT PACK | Akoya Biosciences | FP1497001KT |  |
| Software, Algorithm | FIJI | PMID:29187165 | RRID:SCR_002285 |  |
| Software, Algorithm | CellProfiler | PMID:29969450 | RRID:SCR_007358 |  |
| Software, Algorithm | Prism | Prism 9.3.1 (350) | RRID:SCR_002798 |  |
| Software, Algorithm | RStudio | https://posit.co/ | RRID:SCR_000432 |  |
| Software, Algorithm | Seurat | PMID:29608179 | RRID:SCR_016341 |  |
| Software, Algorithm | ImageJ | https://imagej.net/ | ImageJ<br>RRID:SCR_003070 |  |
| Software, Algorithm | DoubletFinder | PMID:30954475 | RRID:SCR_018771 |  |

**Table S3.**
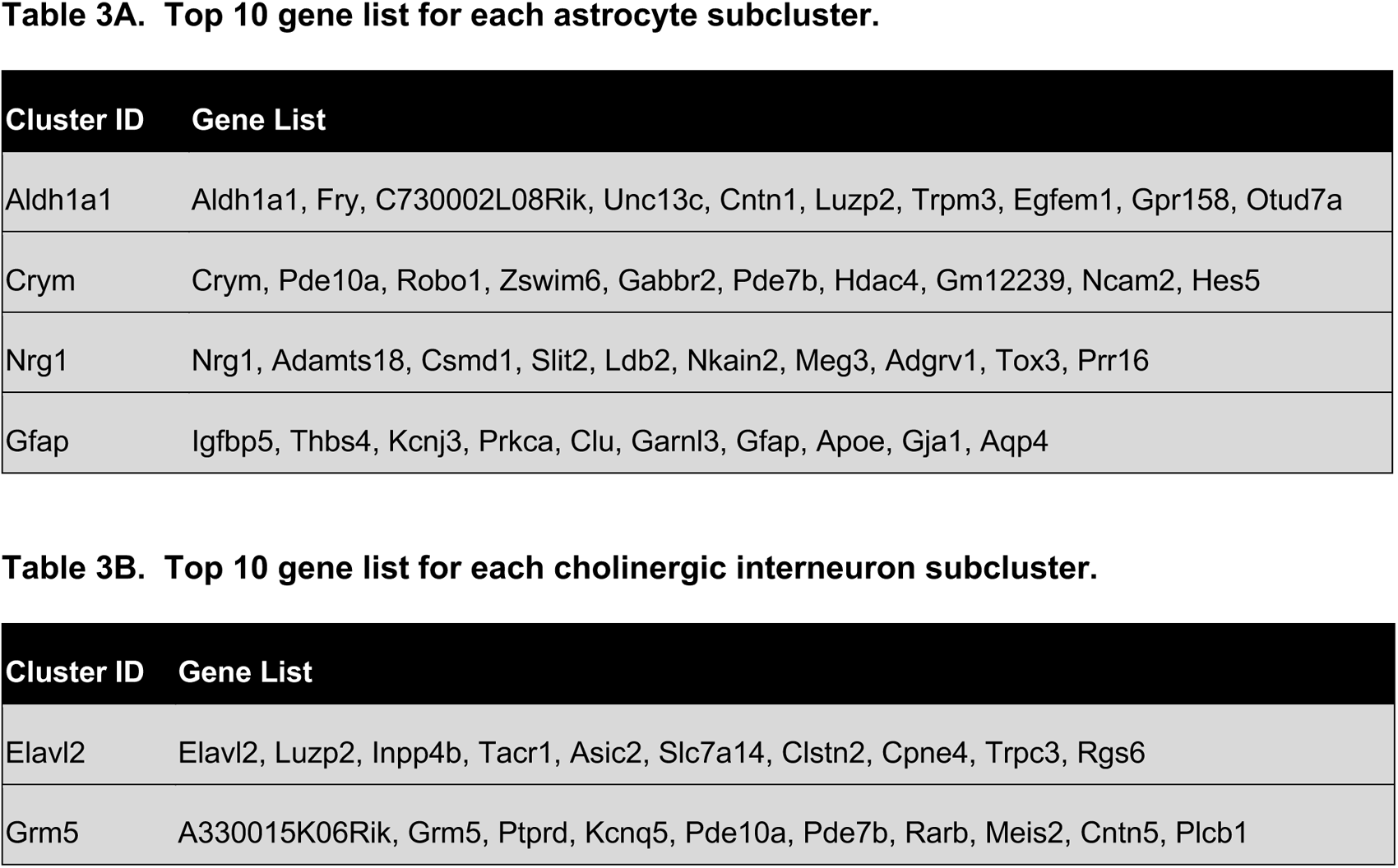
Top 10 gene list for each astrocyte and cholinergic interneuron subcluster.

## Notes

### Competing Interest Statement

The authors have declared no competing interest.

